# Glial vesicular transmitter release is not critical for courtship conditioning memory in *Drosophila melanogaster*

**DOI:** 10.64898/2026.01.10.698790

**Authors:** Paola Virginia Migues, Oliver Hardt

## Abstract

Glial cell function is believed to be critical for the regulation of cognitive processes such as learning and memory but the mechanisms by which glia regulate memory are still poorly elucidated. There is evidence that glial metabolic support and neurotransmitter synaptic clearance are essential for memory processes. But glial cells can also influence neuronal activity by the vesicular release of gliotransmitters in a calcium-dependent manner. Whether gliotransmission also is critical for memory process remains controversial. Here, we explored the role of gliotransmission in *Drosophila* courtship memory using the thermo-sensitive *Shibire^ts^* mutant in combination with the *UAS/Gal4* expression system, in which glial vesicular transmitter release can be rapid and reversibly impaired. We found that the formation, maintenance and decline of courtship memory were unaffected by the blockade of vesicular release of gliotransmitters. Therefore, our data support the view that the release of gliotransmitters is not an ubiquitously necessary regulatory mechanism of memory and forgetting processes.

## Introduction

Glial cells play major supporting roles in the regulation of neuronal functions in the mature nervous system. They control the clearance of neurotransmitters from the synaptic cleft, provide metabolic substrates to adjacent neurons, control cerebral blood flow, participate in homeostatic scaling and potassium buffering, and provide immune function [1, 2]. Besides these metabolic and homeostatic functions glial cells can sense neuronal activity and release transmitters such as glutamate, ATP and d-serine in a calcium-dependent vesicular mechanism involving, like neurons, the SNARE complex [3–7].

Glial cells, in particular astrocytes, are thought to play important roles in neuronal network function, synaptic plasticity and memory [8–13]. An unresolved question is whether the involvement of glial cells in memory processes is mediated solely by their metabolic homeostatic function or also by the calcium-dependent vesicular release of transmitters. Various studies examining the role of glia in memory either have targeted their metabolic and synaptic clearance function or did not dissociate their metabolic from their signaling function [14–18]. Many studies, nevertheless, have used transgenic mice, in which molecules involved in calcium signaling or in vesicle exocytosis were modified, to address how astrocyte signaling contributes to memory and synaptic plasticity. These studies yielded inconsistent results. While some studies have shown that interfering with astrocytic signaling can impair memory and/or synaptic plasticity [6, 7, 19–24], others have found no such effects [21, 25] or detected deficits restricted to certain forms of memory [20, 22–24, 26, 27]. Some of these discrepancies have been attributed to the interventions targeting astrocytic function. For instance, some promoters used for manipulating gene expression selectively in astrocytes could also cause neuronal leakage; furthermore, other interventions, such as protein knockouts, lack precise temporal control and can lead to the engagement of compensatory mechanisms [13]. Some studies have shown that optogenetic and chemogenetic activation of astrocytes modulates memory performance [24, 28, 29]; however, these studies leave unanswered whether the activity of astrocytes modulates memory under physiological conditions.

Therefore, we decided to directly address whether gliotransmission is involved in memory processes by taking advantage of the well-characterized *Shibire^ts^* (*Shi ^ts^*) mutant in *Drosophila melanogaster*. This mutant expresses a temperature-sensitive allele of the *Drosophila* homolog dynamin gene *Shibire*, a molecule critical for synaptic vesicle recycling. The mutant dynamin molecule encoded by *Shi ^ts^* has a single amino acid substitution in the GTPase domain that renders it dysfunctional at temperatures above 29**°**C. As a result, synaptic vesicle recycling is blocked, and, consequently, synaptic vesicles become depleted and transmitter release is supressed [30]. When flies are returned to permissive temperatures, i.e., temperatures below 29**°**C, the molecule reverts to normal function. Thus, transmitter release can be rapidly and reversibly controlled by shifting the ambient temperature to values below or above 29**°**C.

To express *Shi ^ts^* specifically in glial cells we employed the *Gal4/UAS* system. This system consists of a *Gal4* transcriptional activator that is expressed in a specific pattern and a transgene under the control of a *UAS* promoter that is silent in the absence of *Gal4*. To activate the transgene, transcription lines containing the *UAS*-controlled gene are mated to flies expressing the driver *Gal4*. The progeny then expresses the *UAS*-controlled gene in the specific *Gal4* expressing cells [31]. The combination of the *Shi ^ts^* mutation under the control of a *UAS* promoter in conjunction with a glial promoter enhancer driver allows to specifically target glial synaptic vesicle recycling in a precisely timed, reversible, and non-invasive manner. The major advantage of this method for our purpose is that compensatory mechanisms are unlikely to develop, and that the manipulation can be rapidly controlled and therefore restricted to specific memory phases or processes.

In *Drosophila*, as in mammals, glial cells are found in all parts of the nervous system, and their morphology adapts to maximize contact with neurons. They are classified into five different subtypes based on their morphology and location [2, 32]. Here, we target all subtypes, in the whole brain, using the pan-glial driver *Repo-Gal4* [33]. To assess memory performance, we used a courtship conditioning task, a paradigm that has been widely used to study learning and memory processes in *Drosophila* [34–37]. We then examined whether blocking gliotransmission affects the acquisition, maintenance, or the natural decline of memory over time.

## Experimental Procedures

### Fly Strains

The fly strains *Repo-Gal4/TM3* (BDSC #7415), *UAS*-*Shi^ts^(*BDSC #44222), *UAS-EGFP* (BDSC #5431), *Elav-Gal4/CyO* ( BDSC #8765) were obtained from Bloomington Drosophila Stock Center (Indiana University, Bloomington, Indiana, USA). *Canton-S* (*CS*) flies were used for wild-type. *Repo-Gal4* and *UAS-Shi^ts^* were outcrossed to the *CS* background flies for six generations to harmonize their genetic background.

Flies were kept on a 12:12 h light-dark cycle inside an insect incubator (PHCbi MIR-254) at 22**°**C and 65-75% humidity, except for inducing the *Shibire^ts^* dysfunction, when the temperature was increased to 30**°**C. Flies were kept in 175 ml plastic vials containing 50 ml of food and a folded filter paper (Whatman 595 ½) to increase the pupation area. The food used for stock maintenance and for the production of males for the behavioural task contained 0.5% agar (Genesee Scientific), 2.75% active yeast (Red Star), 5.2% corn flour (Genesee Scientific), 11% sugar (Fisher Scientific), 0.05% methylparaben (Genesee Scientific), and 0.5% propionic acid (Sigma Aldrich) in distilled water [35]. Only for the production of mated females or for housing the male test subjects in the courtship task we used a specialised food (“power food”), which leads to a large number of flies, containing: 0.8% agar, 8% yeast 2% yeast extract (MP Biomedical), 2% peptone (Sigma Aldrich), 3% sucrose, 6%, glucose (Sigma Aldrich), 0.05% MgSO4 (Sigma Aldrich), and 0.05% CaCl_2_ (Merck), 0.05% methylparaben, and 0.5% propionic acid in distilled water [35].

### Behavioural task

Flies were trained and tested using a courtship conditioning task and apparatus as described in Koemans et al. [35]. In this task virgin male flies are exposed to a previously mated females. Male flies court the females in a set of stereotyped courtship behaviors but as the females are not receptive, they reject the mating attempts. Thus, males subsequently suppress their courtship behavior towards premated females. The courtship suppression can be evaluated in a testing session by measuring the time the male flies spend courting a mated female.

### Culture preparation

Fourteen days before the experiment we started the cultures to produce male test subjects using 4–5 d old flies*. Repo-Gal4/UAS-Shi^ts^* were generated by crossing virgin *Repo-Gal4* females with *UAS-Shi^ts^* males. For control groups virgin *Repo-Gal4* females were crossed with male wild type *CS* (*Repo-Gal4/CS*), and virgin *CS* females with males *UAS-Shi^ts^* (*CS/UAS-Shi^ts^*). Approximately 40 females and 30 males were placed in each vial. To produce mated females, we used 4-5 d old *CS* flies. All cultures were transferred to a new vial every 24 h over 5 days.

### Collection of male subjects and mated female flies

Male flies of the correct genotype were collected within 30 min after eclosion using carbon dioxide anaesthesia at a flow rate of 2 L/min. Each individual fly was transferred using an aspirator to an individual well of a 96-well housing block (Quiagen) containing 500 µl of power food where they remained for four days. For the preparation of mated females newly eclosed females and males *CS* flies were collected under anaesthesia and placed in a mating vial containing power food and a small amount of yeast paste. Approximately 20 females and 10 males were placed in each vial.

### Training

Five days after eclosion we collected anaesthetized mated females from the mating vial and placed them in an individual well in a 96-well block. Then we transferred an individual 4-day-old male without anesthesia with an aspirator from the housing block to a well containing a premated female. For the naïve controls males were transferred to a well without female. The pairs were left at 22**°**C or 30**°**C for 8 h in experiments testing long-term memory or 1 h in experiments testing short-term memory. At the end of training males or naïve males were transferred without anaesthesia to an individual well in a new housing block and left at 22**°**C or 30**°**C until testing. This protocol was repeated over 4 d to achieve group sizes of approximately 40 flies.

### Test

The test was conducted in a specialised courtship chamber containing 18 arenas (10 mm diameter 4 mm deep) each with a divider in the middle (Fig.1 A). The chambers were based on the design of Koemans et al. [35] and printed at McGill Physics Makerspace and 3D Print Lab. All dividers of the arenas can be opened simultaneously, so that courting can begin in all arenas at the same moment. Males were placed individually with an aspirator and without the use of anaesthesia into each arena with the dividers closed. One mated female (5 d old) was transferred to the other half of each arena containing a male. Then the dividers were opened to allow simultaneous contact between all paired flies. Behaviour was recorded with a video camera (Sony, HDR-CX405) for 10 min.

**Figure 1.**
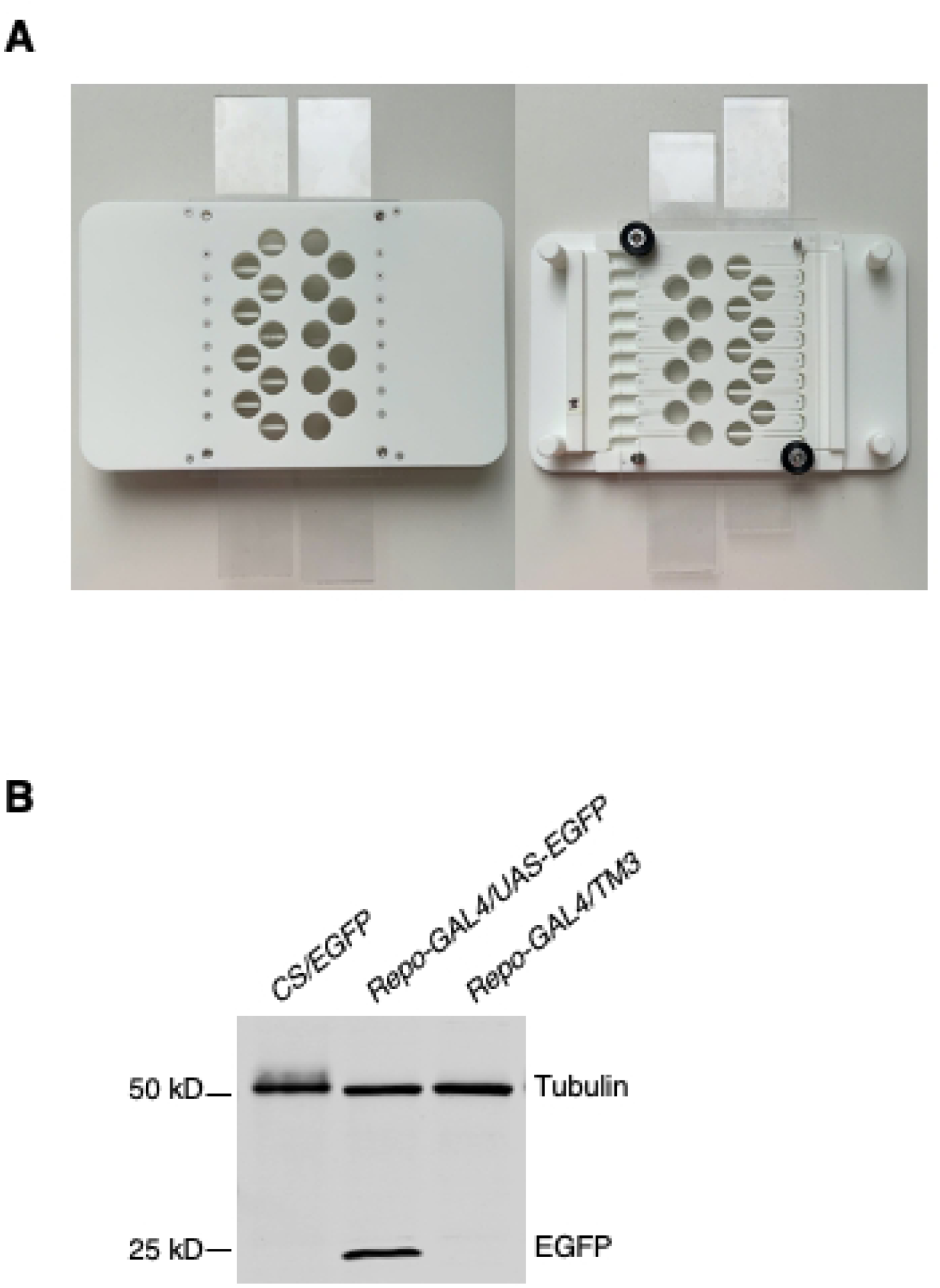
**A**. Courtship chamber used for the courtship conditioning test. The top side of the chamber is shown on the left picture and the bottom side on the right. The dividers are closed when placing females and males in each side of the arena and they are open simultaneously to allow the interaction of males and females. **B**. Efficacy validation of the driver *Repo-Gal4*. Representative Western blot against EGFP and tubulin performed on fly head homogenates of *Repo-Gal4/ UAS-EGFP*, *Repo-Gal4/ TM3* and *CS/UAS-EGFP* flies. EGFP expression was detected only in *Repo-Gal4/ UAS-EGFP* brains. Each sample contains the homogenate of 40 fly heads.

### Data analysis

Courtship behavior was defined as the male following the female, vibrating the wings, licking, or attempting copulation. We manually determined the time flies spent expressing courtship behaviour during test, using in-house software to score the recorded videos. A courtship index (CI) was calculated by dividing the time the male engaged in courtship behaviour by the 10 min of the testing period. Memory is reflected in a decrease of the CI of the trained versus the naïve, untrained control [35].

Data were tested for normality (Shapiro-Wilk). Because the data did not follow a normal distribution, groups were compared with the non-parametric Wilcoxon rank-sum test using JMP (JMP Statistical Discovery) statistical software.

### Immunoblotting

Frozen 4-5 d old flies (kept at -80°C) were decapitated by vigorous vortexing for 30 s; the heads (40 heads per sample) were collected under a dissecting microscope and homogenized in 40 µl of RIPA buffer (Pierce, Thermo Scientific 25 mM Tris-HCl pH 7.6, 150 mM NaCl, 1% NP-40, 1% sodium deoxycholate, 0.1% SDS) containing Halt^TM^ protease inhibitor cocktail (Thermo Scientific) and EDTA 5 mM. They were centrifuged at 16000 x g at 4°C for 20 min. The supernatant was collected and mixed with equal volume of 2X SDS sample buffer (63 mM Tris HCl, 10% glycerol, 2% SDS, 0.0025% bromophenol blue, pH 6.8) and boiled at 85°C for 10 min. After boiling DTT was added to each sample at a final concentration of 0.1 M.

Samples (10 µl) were loaded in a 4-20% Tris-Glycine gel (Novex Wedge Well, Invitrogen). The gels were run in an electrophoresis chamber (X-cell Surelock^TM^ minigel, Invitrogen) at 120 V constant voltage with Tris-Glycine SDS Running buffer (25 mM Tris, 192 mM glycine, 0.1% SDS, pH 8.3), and then transferred onto a 0.45 micrometer nitrocellulose membrane (BioRad) at a 25 V constant voltage for 60 min in Tris-Glycine transfer buffer (12 mM Tris, 96 mM Glycine, 20% Methanol). After washing with TBS, the membranes were blocked with blocking solution (Intercept Li-Cor). The membranes were then incubated simultaneously with an antibody against GFP (1/2000, Abcam AB6556) and an antibody against tubulin (1/2000, Abcam AB44928) in blocking solution containing 0.2% Tween for 1 h. After washing four times with TBS-tween (0.2% Tween) they were incubated with fluorescent secondary antibodies (1/20000, IRDye 800, iRDye690, Lycorin) in TBS-Tween for 1 h. The membranes were rinsed four times in TBS-Tween with a final rinse in TBS and scanned wet with a Li-COR Odyssey Dlx scanner.

## Results

### Efficacy validation of *Shibire^ts^* and the driver Repo-Gal4

To disrupt transmitter release we used the thermo-sensitive *Shibire* mutant strain *UAS-Shi^ts^* described above. When *UAS-Shi^ts^* is combined with a ubiquitous neuronal enhancer driver such as *Elav-Gal4,* a shift in temperature to 30°C results in blocked synaptic transmission and thus paralysis [30]. When combined with a specific glial promoter enhancer *Repo-Gal4* the blockade of vesicle trafficking occurs only in glial cells and thus does not cause paralysis [33].

To confirm the efficacy of *Shi ^ts^* mutation in our *UAS-Shi^ts^* flies we generated *Elav-Gal4/UAS-Shi^ts^* flies by crossing *UAS-Shi^ts^* males flies with virgin *Elav-Gal4* females and observed their behavior when placed at 30°C. The *Elav-Gal4/UAS-Shi^ts^* flies became motionless at 30°C whereas *Elav-Gal4/CS* flies or *Repo-GAL4*/ *UAS-Shi^ts^* exhibited normal motor activity (S1 Video). When *Elav-Gal4/UAS-Shi^ts^* flies were returned to 22**°**C they regained normal locomotor activity. These findings confirm the efficacy and reversibility of dynamin disfunction at 30**°**C in the flies used in our behavioural studies. Additionally, they show that the *Repo-Gal4* driver does not result in expression of *Shi ^ts^* in neurons.

We also assessed the efficacy of the *Repo-Gal4* driver by crossing *Repo-Gal4* females flies with *UAS-EGFP* males. These flies express the enhanced green fluorescence protein (EGFP) under the control of a *UAS* promoter. As control we used *CS/UAS-EGFP and Repo-Gal4/TM3*. The expression of EGFP was determined by Immunoblotting. As expected, we detected the presence of EGFP only in *Repo-Gal4/UAS-EGFP* (Fig.1B).

### Gliotransmission blockade does not affect learning

To assess memory performance, we used a courtship conditioning task as described in the Method section. We first explored if blocking gliotransmission during the training session would affect the formation of long-term memory. *Repo-Gal4/UAS-Shi ^ts^* flies were trained at 30**°**C to induce dysfunction of the glial dynamin molecule expressed by *Shi^ts^*. Control naïve flies were handled and exposed to the same conditions but without being exposed to a female. For genotype control we used *Repo-Gal4/CS* and *CS*/*UAS-Shi ^ts^* flies. For the control of the induction of the dynamin dysfunction and of the effect of temperature on behavior, all three genotypes were also trained at 22**°**C, a temperature at which dynamin function is undisturbed. After the 8 h training session all flies were housed at 22**°**C and then tested at 22**°**C, 22 h after the end of training. Trained flies from all genotype groups had statistically significantly reduced CI compared to their matched naïve controls either at 22**°**C (*Repo-Gal4/UAS-Shi ^ts^ p*<0.0001; *Repo-Gal4/CS p*<0.0005; *CS*/*UAS-Shi ^ts^ p*=0.0092 Wilcoxon rank-sum test) or at 30**°**C (*Repo-Gal4/UAS-Shi ^ts^ p*<0.0005; *Repo-Gal4/CS p*<0.0005; *CS*/*UAS-Shi ^ts^ p*=0.0001 Wilcoxon rank-sum test; Fig. 2A), indicating intact courtship memory. These results show that the blockade of gliotransmission during training had no effect on the formation of long-term memory.

**Figure 2.**
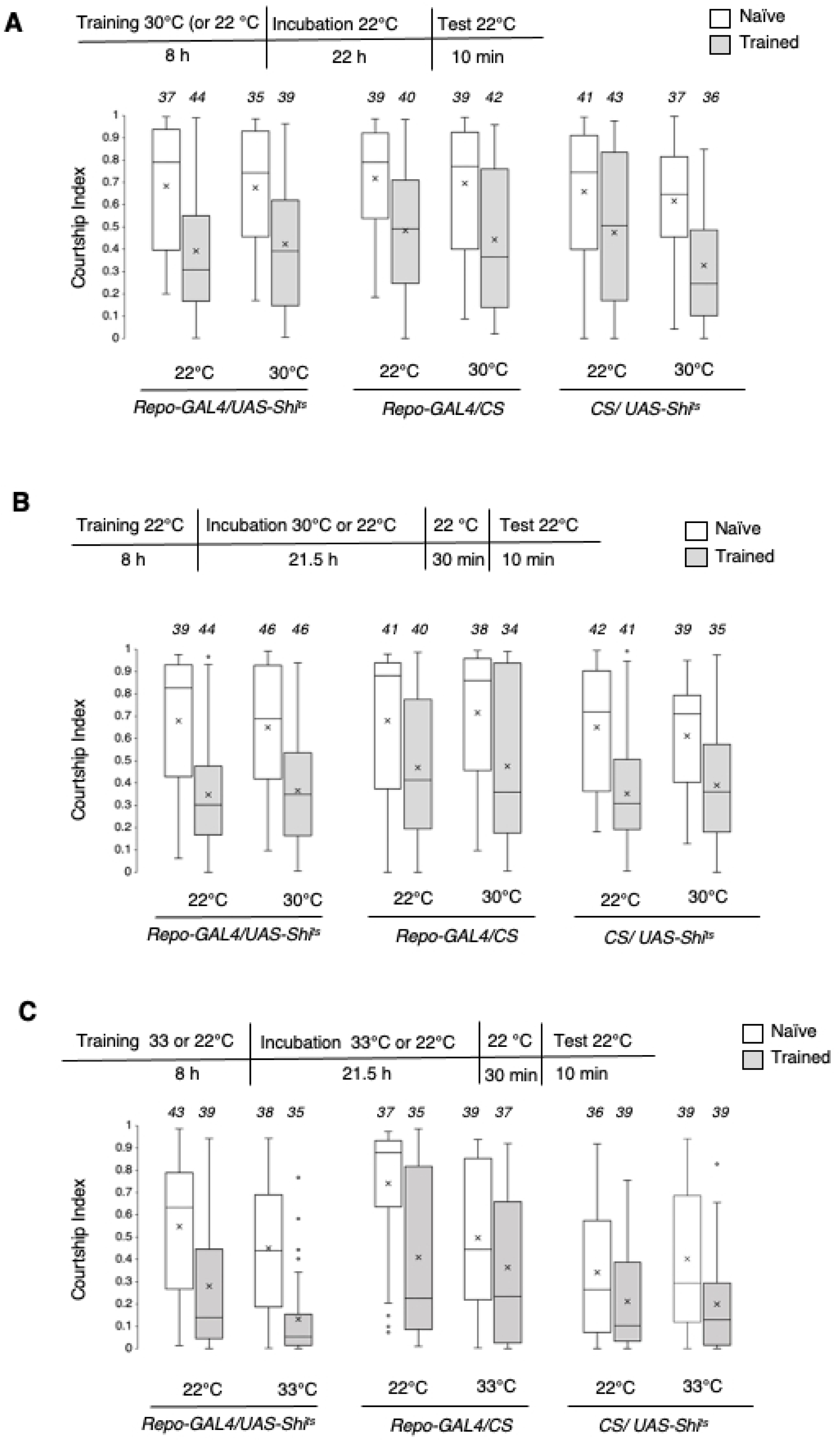
Gliotransmission blockade does neither affect the acquisition nor the maintenance of memory. Box plots including outliers and mean values (x) comparing the distribution of CI values of naïve and trained flies of *Repo-Gal4/UAS-Shi ^ts^*, *Repo-Gal4/CS and CS*/*UAS-Shi ^ts^* genotypes. Blocking glial vesicle transmitter release during training (**A, C**) or after training (**B, C**) at 30°C (**A, B**) or 33°C (**C**) did not affect memory performance at 22 h later. Trained flies from all genotypes had statistically significant lower CI than the corresponding naïve group (*p*<0.01, Wilcoxon rank-sum test) indicating intact memory at both permissive (22**°**C) or *Shi* dysfunction (30**°**C or 33**°**C) conditions. The numbers in italics above each box plot indicate the sample size (*n)* for the corresponding group.

### Gliotransmission blockade does not affect memory maintenance

We performed an experiment similar to the one above, but trained animals at 22**°**C and then immediately thereafter we housed the flies at 30**°**C for 21.5 h to impair glial dynamin function after learning. Thirty min before the test flies were transferred to 22**°**C to reverse the impairment, and the test was also performed at 22**°**C. All groups exhibit courtship memory as indicated by reduced CI in trained flies compared to their naïve controls at both 22**°**C (*Repo-Gal4/UAS-Shi ^ts^ p*<0.0001; *Repo-Gal4/CS p*=0.0075; *CS*/*UAS-Shi ^ts^ p*<0.0001 Wilcoxon rank-sum test) and at 30**°**C (*Repo-Gal4/UAS-Shi ^ts^ p*<0.0001*; Repo-Gal4/CS p*=0.0230; *CS*/*UAS*-*Shi ^ts^ p*=0.0006 Wilcoxon rank-sum test; Fig. 2B). These results indicate that suppression of vesicular gliotransmission after training does not affect the maintenance of memory.

A previous study has shown that exposing flies expressing *UAS-Shi^ts^* in cortex glia (a *Drosophila* glia subtype) to 33**°**C for 6 h following training impairs long-term associative olfactory memory [38]. Therefore, to rule out that the lack of effect we observed in our study was not due to the different restrictive temperature we used, we repeated the above experiments at 33**°**C. We found that long-term memory was unaffected when *Repo-Gal4/UAS-Shi^ts^* flies were exposed to 33**°**C during the 8 h training period and for 21 h afterward. All genotypes exhibited memory at both 22**°**C (*Repo-Gal4/UAS-Shi ^ts^ p*=0.0002, *Repo-Gal4/CS p*=0.0002, *CS*/*Shi ^ts^* p=0.0456 Wilcoxon rank-sum test) and 33**°**C (*Repo-Gal4/UAS-Shi ^ts^ p*<0.0001; *Repo-Gal4/CS p*=0.0322; *CS*/*UAS-Shi ^ts^* p=0.0035 Wilcoxon rank-sum test; Fig. 2C).

### Gliotransmission blockade does not affect short-term memory

We next tested the effects of gliotransmitter release on short-term memory. Flies were trained for 1 h at 22**°**C and tested 3 h later at 22**°**C. The impairment of dynamin was induced immediately after training and reversed 30 min before the test. All genotypes showed intact memory at both 22**°**C (*Repo-Gal4/UAS-Shi ^ts^ p*=0.0194; *Repo-Gal4/CS p*=0.0020; *CS*/*UAS-Shi ^ts^ p*=0.0002 Wilcoxon rank-sum test) and at 30**°**C (*Repo-Gal4/UAS-Shi ^ts^ p*<0.0001; *Repo-Gal4/CS p*=0.0015; *CS*/*UAS-Shi ^ts^ p*=0.0216 Wilcoxon rank-sum test; Fig. 3A) indicating that disruption of gliotransmitter release has no effect on short-term memory retention.

**Figure 3.**
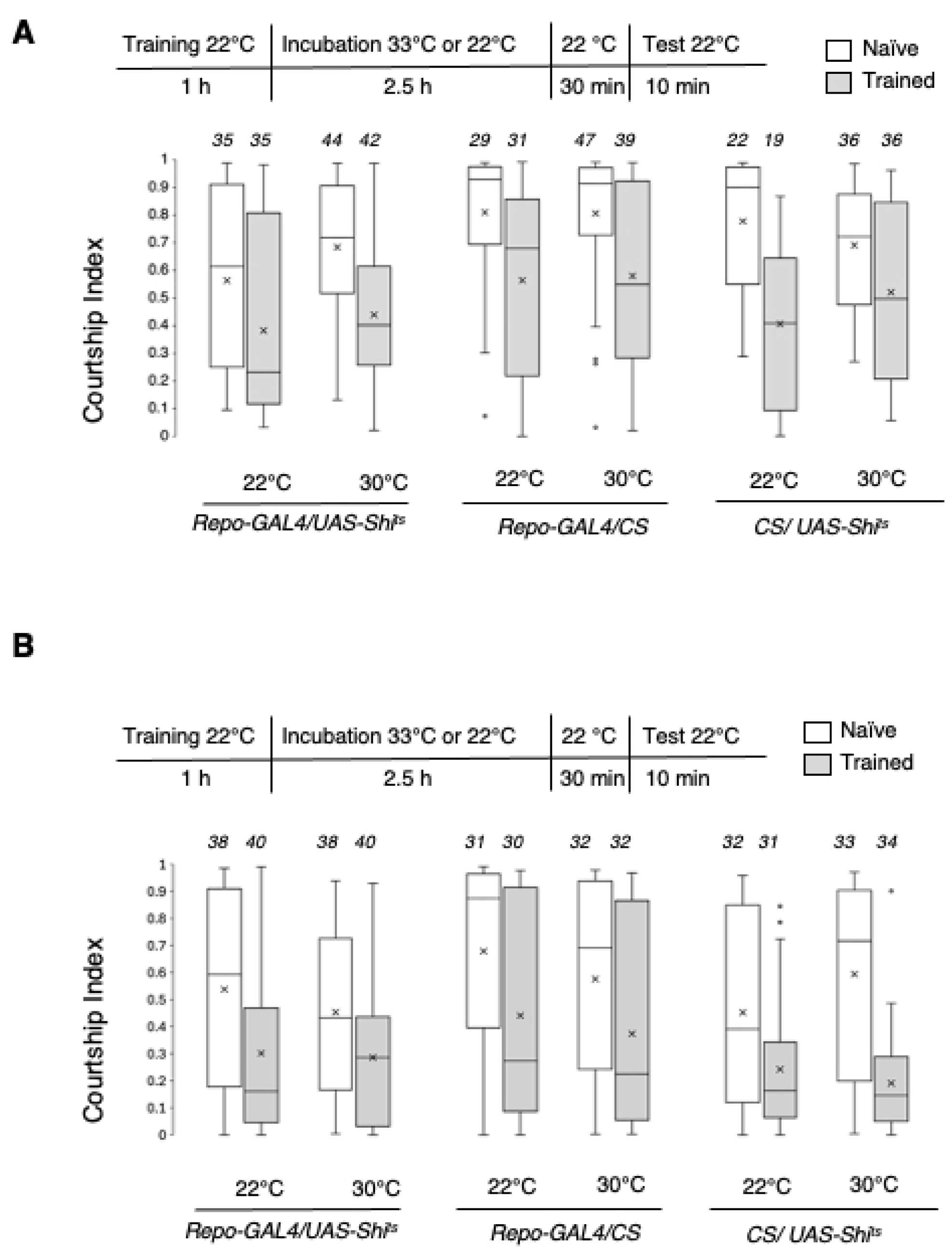
Gliotransmission blockade does not affect short-term memory. Box plots including outliers and mean values (x) comparing the distribution of CI values of naïve and trained flies of *Repo-Gal4/UAS-Shi ^ts^*, *Repo-Gal4/CS and CS*/*UAS*-*Shi ^ts^* genotypes. Blocking glial vesicle transmitter release 1 h after training at 30**°**C (**A**) or 33**°**C (**B**) did not affect memory performance at 3 h later. Trained flies from all genotypes had statistically significant lower CI than the corresponding naïve group (*p*<0.01, Wilcoxon rank-sum test) indicating intact memory at both permissive (22**°**C) or *Shi* dysfunction (30**°**C or 33**°**C) conditions. The numbers in italics above each box plot indicate the sample size (*n)* for the corresponding group.

As we did for long-term memory, we repeated this experiment with a temperature shift to 33**°**C instead of 30**°**C. We obtained results similar to those observed with the 30**°**C temperature change. The CI was significantly lower in the trained groups than in the naïve controls at both 22**°**C: *Repo-Gal4/UAS-Shi ^ts^ p*=0.0013; *Repo-Gal4/CS p*=0.0078; *CS*/*UAS*-*Shi ^ts^ p*=0.0097) and 33**°**C (*Repo-Gal4/UAS-Shi ^ts^ p*=0.0025; *Repo-Gal4/CS p*=0.0291; *CS*/*UAS*-*Shi ^ts^ p*<0.0001 Wilcoxon rank-sum test; Fig 3B).

### Gliotransmission blockade does not affect forgetting

The results above suggest that disruption of synaptic vesicle trafficking in glial cells it is neither detrimental for the formation nor for the maintenance of memory. We therefore next explored if gliotransmission blockade could prevent the natural decline of memory. We first determined when flies no longer express courtship conditioning memory after training at a permissive temperature (22**°**C). We trained Repo*-Gal4/UAS-Shi ^ts^* flies for 8 h and tested them after 1, 3, 5 or 8 d. We found that trained flies had significant lower CI versus naïve flies after 1 d (*p*<0.0001) and 3 d (*p*=0.0021), but not after 5 d (*p*=0.1464) or 8 d (*p*=0.6086 Wilcoxon rank-sum test; Fig. 4A). These findings indicate that memory starts to decline 3 d after training and is no longer expressed after 5 d.

**Figure 4.**
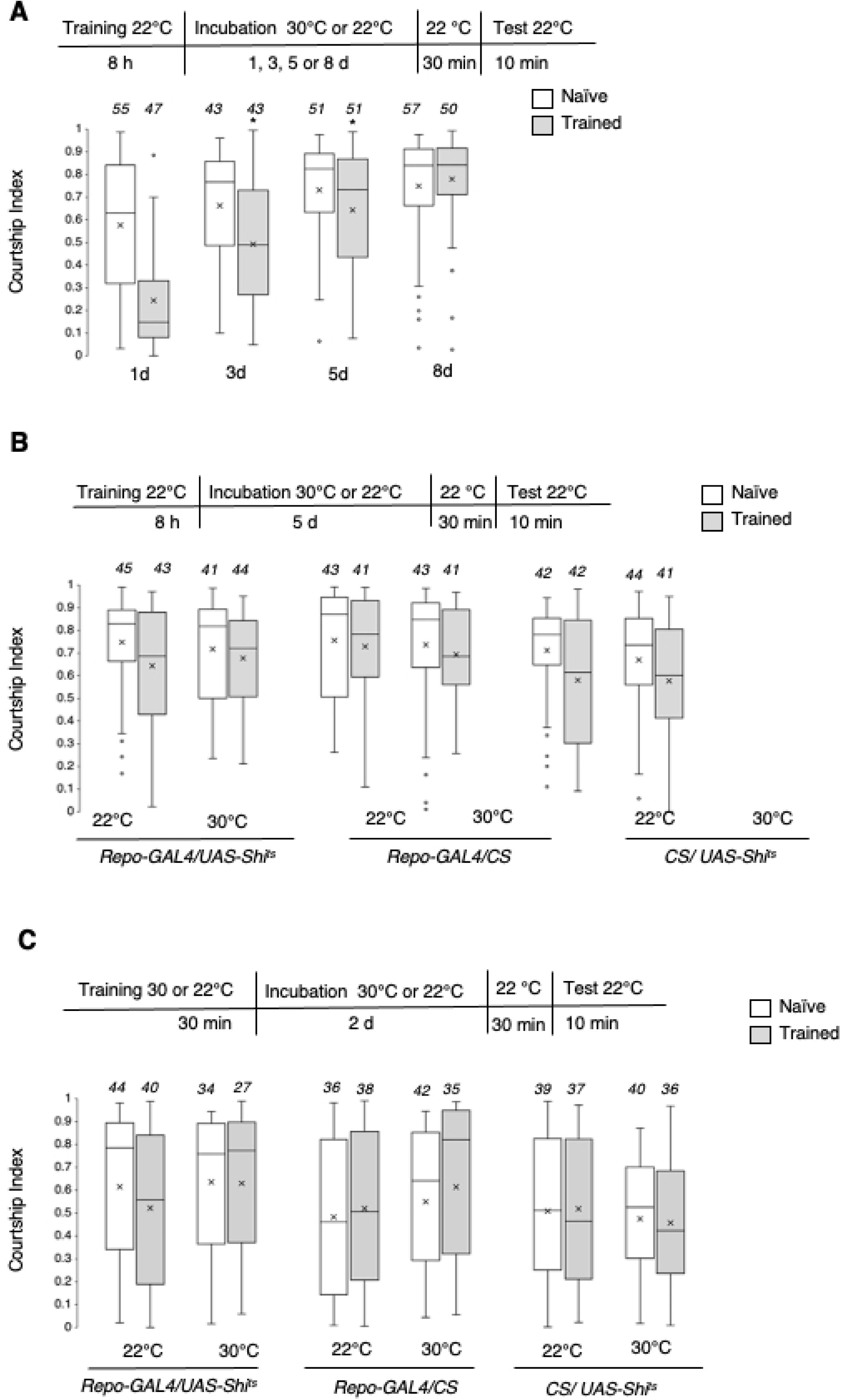
Gliotransmission blockade does not affect forgetting. **A.** Box plots including outliers and mean values (x) comparing the distribution of CI values of naïve and trained flies of *Repo-Gal4/UAS-Shi ^ts^* without induction of *Shi* dysfunction at different times after 8 h training. Flies tested after 1 d or 3 d show statistically lower CI than their respective naïve control group (*p*<0.01, Wilcoxon rank-sum test). No statistical differences were detected between trained and naïve flies when tested at 5 d or 8 d after training. **B, C**. Box plots including outliers and mean values (x) comparing the distribution of CI values of naïve and trained flies of *Repo-Gal4/UAS-Shi ^ts^*, *Repo-Gal4/CS and* CS/*UAS*-*Shi ^ts^* genotypes. Blocking glial vesicle transmitter release after 8 h training did not prevent memory loss 5 d later (**B**). Blocking glial vesicle transmitter release during and after 1 h training did not prevent memory loss 2 d later (**C**). No significant differences were detected between trained and naïve flies either at permissive (22**°**C) or *Shi* dysfunction (30**°**C) conditions (**B, C**). The numbers in italics above each box plot indicate the sample size (*n)* for the corresponding group.

To explore whether the disruption of synaptic vesicle transmitter release in glial cells could prevent memory loss, we assessed memory performance 5 d after the end of an 8 h training session. We shifted the temperature to 30**°**C immediately after training to disrupt dynamin function and lowered the temperature to 22**°**C 1 h before the test. There was no significant difference between trained and naïve flies housed at 22**°**C (*Repo-Gal4/UAS-Shi ^ts^ p*=0.1342, *Repo-Gal4/CS p*=0.4078, *CS*/*UAS-Shi ^ts^ p*=0.0997 Wilcoxon rank-sum test) indicating the absence of courtship memory. Similarly, the CI of trained flies housed at 30**°**C was also not significantly different from the CI of naïve flies (*Repo-Gal4/UAS-Shi ^ts^ p*=0.1728; *Repo-Gal4/CS p*=0.0707; *CS*/*UAS*-*Shi ^ts^ p*=0.0616 Wilcoxon rank-sum test; Fig. 4B). These findings indicate that the blockade of gliotransmission does not affect forgetting.

We then assessed the effect of gliotransmission blockade on forgetting using a weak 30 min training protocol and tested memory two days after training. The temperature was shifted to 30**°**C 30 min before training and maintained until 30 min before the test performed 2 d later. There was no difference in courtship behaviour between trained and naïve flies at either 22**°**C (*Repo-Gal4/UAS-Shi ^ts^ p*=0.1720; *Repo-Gal4/CS p*=0.6303; *CS*/*UAS*-*Shi ^ts^ p*=0.8720 Wilcoxon rank-sum test) or 30**°**C (*Repo-Gal4/UAS-Shi ^ts^ p*=0.8107; *Repo-Gal4/CS p*=0.1449; *CS*/*UAS-Shi ^ts^ p*=0.6584 Wilcoxon rank-sum test; Fig. 4C). These results indicate that blocking gliotransmission does not prevent the decline of memory induced by weak training, consistent with the results obtained with the 8 h training.

## Discussion

Our study demonstrates that normal learning, memory and forgetting can occur in the absence of vesicular transmitter release from glial cells in *Drosophila*. It is highly unlikely that the lack of alterations in memory performance are due to the engagement of potential compensatory mechanism because we used a very rapid and reversible induction method that allows the transmitter release suppression to occur within minutes of induction. We cannot rule out, however, that alternative ‘back up’ mechanisms that may be already in place could get engaged when dynamin is disrupted for a long time, but this seems unlikely for the short-term memory test. Thus, gliotransmission appears not to be critical for short-term or long-term courtship conditioning memory.

Our findings seem to be in contrast with the results of a previous study showing that blocking vesicle release in cortex glia, a *Drosophila* glia subtype, impairs long-term associative olfactory memory [38]. Importantly, this study also reports that the memory impairment following gliotransmission blockade depends on the training protocol used; specifically, gliotransmission seems only required when flies are subjected to spaced training sessions, but not when massed training is used. Thus, taken together, it is possible that in *Drosophila* the involvement of glia vesicle release may be limited to specific learning regimes and types of memory. A similar pattern has been observed in mice where several studies have shown that gliotransmission is required only in some memory tasks [23, 24, 26], and only for remote, but not recent memories [20, 22, 23].

Several studies in mice have cast doubt on the notion that physiological calcium-dependent release of gliotransmitters is a major and ubiquitous modulator of memory processes. This is because intact synaptic plasticity and memory performance in transgenic mice occurs normally when gliotransmission is impaired [21, 25]. Additionally as mentioned above several studies have found that gliotransmission does not seem to be required for recent memories. Our findings are in line with this idea and provide evidence extending this view to *Drosophila*.

## Acknowledgments

We thank Dr Thomas Brunner, Robert Turner, and Dr Xiao Shang from the McGill Physics Makerspace and 3D Print Lab (https://makerspace.physics.mcgill.ca/index.html) for constructing the courtship chambers.

---

**S1.** Efficacy validation of *Shibire^ts^*. Video showing *Elav-Gal4/UAS-shi^t^*, *Repo-Gal4/UAS-Shi ^ts^* and *Elav-Gal4/CS* flies at 22°C and 30°C. Only *Elav-Gal4/UAS-Shi^t^* became motionless at 30°C.

